# Re-evaluating the Need for Double Centrifugation in Plasma Cell-Free DNA Analysis

**DOI:** 10.64898/2026.01.30.702926

**Authors:** Y Wang, PA Shaw, A Vallon, Naief L Tavares, A Hicks, M Ednie, L Ritzert, FRG Amrit, T Chu, CG McKennan, DG Peters

## Abstract

Plasma cell-free DNA (cfDNA) is a central analyte in liquid biopsy applications spanning prenatal testing, oncology, and epigenomic profiling. To minimize contamination by high-molecular-weight genomic DNA (gDNA) released from nucleated blood cells, standard pre-analytical workflows typically mandate a double-centrifugation protocol prior to cfDNA extraction. This requirement has limited the use of many existing plasma biorepositories that were prepared using only a single low-speed centrifugation step. In this study, we evaluated whether single-spun plasma is sufficient for accurate cfDNA analysis when samples are processed under controlled conditions. Using paired single- and double-spun plasma aliquots derived from the same early-pregnancy maternal blood samples collected in EDTA tubes, we performed whole-genome DNA methylation sequencing and assessed cfDNA integrity across multiple orthogonal dimensions. These included cell-type proportion deconvolution using large and small DNA methylation reference signatures, CpG-level methylation rate estimation with explicit variance modeling, beta-binomial-corrected correlation analyses across libraries, cfDNA fragment length profiling, and genotype-based fetal fraction estimation. Across all analyses, we found no evidence that a second high-speed centrifugation step improved accuracy, reduced technical variability, or enhanced analytical fidelity. Cell-type proportion estimates and CpG-level methylation rates were statistically indistinguishable between single- and double-spun plasma, fragment length distributions were nearly identical, and fetal fraction estimates showed near-perfect concordance. Together, these results demonstrate that a single low-speed centrifugation step is sufficient for high-fidelity cfDNA methylation, fragmentomic, and genotyping analyses. Our findings support the expanded use of legacy single-spun plasma collections for liquid biopsy research and assay development and motivate a re-evaluation of rigid double-centrifugation requirements in cfDNA workflows.

## INTRODUCTION

Liquid biopsy has emerged as a transformative approach in molecular diagnostics, enabling the non-invasive detection and monitoring of a wide range of physiological and pathological conditions through the analysis of circulating nucleic acids (Wan et al., 2017, Merker et al., 2018). Among these, cell-free DNA (cfDNA) has become a key analyte in applications including prenatal testing, oncology, organ transplantation, and infectious disease surveillance (Chu et al., 2015, Bettegowda et al., 2014, De Vlaminck et al., 2015). cfDNA in plasma consists primarily of short, nucleosome-protected DNA fragments that originate from apoptotic and necrotic cells (Jahr et al., 2001, Snyder et al., 2016).

The low abundance of cfDNA, its highly fragmented nature, and susceptibility to contamination by genomic DNA (gDNA) from lysed leukocytes necessitate rigorous pre-analytical processing protocols to ensure analytical validity (El Messaoudi et al., 2013, Devonshire et al., 2014). A critical component of these protocols is the double centrifugation of plasma prior to cfDNA extraction (Chiu et al., 2001). The first centrifugation, typically performed at 1,600–2,000 × *g* for 10 to 15 minutes at 4°C, serves to separate plasma from the cellular components of blood (Chan et al., 2005, Diehl et al., 2008). However, despite this initial step, residual leukocytes and platelets can remain in the supernatant, particularly if the interface with the buffy coat is disturbed. These remaining cells pose a substantial risk for *ex vivo* lysis during sample handling, leading to the release of long fragment gDNA that dilutes the cfDNA population and undermines the specificity of downstream analyses.

To minimize the problem of *ex vivo* cell lysis, a second high-speed centrifugation, often at 16,000 × *g* for 10 minutes, is routinely employed to clarify the plasma by removing cellular debris and apoptotic bodies that may contain contaminating DNA (Kang et al., 2016). This double-spin strategy for plasma processing has been shown to significantly improve cfDNA purity and consistency across diverse applications. For example, in cancer liquid biopsy, even trace amounts of gDNA contamination can obscure the detection of low-allele frequency tumor mutations, leading to false-negative or false-positive variant calls (El Messaoudi et al., 2013). Similarly, in prenatal testing, contamination with maternal gDNA can reduce the fetal fraction and compromise assay sensitivity (Chiu et al., 2001).

Numerous studies have highlighted the detrimental effects of delayed processing or inadequate centrifugation on cfDNA quality. Unprocessed EDTA blood samples stored at room temperature for more than six hours can exhibit a marked increase in total DNA concentration due to leukocyte lysis (Chan et al., 2005). To mitigate these effects, blood collection tubes containing stabilizing agents, such as Streck Cell-Free DNA BCTs, have been developed to preserve white blood cell integrity and stabilize cfDNA profiles for up to several days. However, even when such tubes are used, a second centrifugation step remains essential to ensure optimal sample clarity and to prevent carryover of non-cell-free material.

Unfortunately, the reliance on “double-spun” plasma as a source of cfDNA generally requires that blood samples be collected and processed prospectively using the above guidelines. This is because most existing plasma biorepositories, especially those that predate recent increasing interest in liquid biopsy, have been assembled using plasma that has been centrifuged only once at low speed (usually ∼1500 x g). Therefore, based upon the conventional wisdom outlined above, these samples are not considered to be generally useful for liquid biopsy analysis of cfDNA. This is a major limitation to progress because of the high cost and time burden of prospectively recruiting double-spun plasma from human cohorts of interest. This problem is compounded further by the fact that interest in liquid biopsy of cfDNA is expanding rapidly across a variety of clinical realms, driven to considerable extent by interest in the epigenomic information conveyed and its utility both for foundational research and the development of screening/diagnostic tools for clinical use. Therefore, in this study, we tested the hypothesis that single-spun plasma recovered from blood collected in purple top EDTA tubes has utility for liquid biopsy of cfDNA and, importantly, whether it may have equivalent utility to that of its double-spun counterpart.

Despite the widespread adoption of double centrifugation protocols for plasma processing, the analytical benefit of an additional high-speed centrifugation step has rarely been evaluated using modern, genome-wide cfDNA assays. In particular, it remains unclear whether double centrifugation measurably improves analytical fidelity for epigenomic, fragmentomic, or allele-specific cfDNA analyses when plasma is initially separated using a single low-speed spin under controlled conditions.

To our knowledge, no prior study has directly evaluated, using paired plasma aliquots from the same blood draw, whether a second high-speed centrifugation step measurably improves analytical fidelity for genome-wide cfDNA methylation, cell-type deconvolution, fragmentomic analysis, and genotype-based fetal fraction estimation. This study provides a comprehensive, quantitative assessment of a long-standing pre-analytical assumption in liquid biopsy workflows.

## METHODS

### Human Subjects

Maternal peripheral blood samples, collected between 10- and 13-weeks’ gestation, were obtained from three pregnancies. Samples were centrifuged for 10 minutes at 1,500 x g at 4°C and then 50% was removed and this aliquot centrifuged at 15,000 x g at 4°C for further 10 minutes. Plasma samples were immediately frozen in 1mL aliquots and stored at −80°C prior to DNA extraction and sequencing library construction.

### Whole Genome DNA Methylation Sequencing of Plasma Cell-Free DNA

Extracted plasma cell-free DNA samples were converted into Illumina-compatible methylation libraries using NEBNext Enzymatic Methyl-seq Kit (New England Biolabs, Ipswich, MA, USA). Final libraries were quantified on Qubit Flex Fluorometer using Qubit 1x dsDNA High Sensitivity assay (Thermo Fisher Scientific, Waltham, MA, USA), assessed for fragment distribution and quality by 2100 Bioanalyzer High Sensitivity DNA assay (Agilent, Santa Clara, CA, USA), and sequenced on NovaSeq X.

### Cell Type Proportion Analysis

We sought to evaluate whether the number of centrifugation steps had an impact on the accuracy of cell type proportion estimates. We estimated the proportions of 48 cell types using the standard binomial deconvolution model (see equation 2 in Caggioano et al. (Caggiano et al., 2021) and a “large” (Figure 1) or “small” (Figure S1) DNAm signature containing 6,778,063 and 78,536 CpGs that we derived from Loyfer et al. (Loyfer et al., 2023). We considered two signatures to ensure conclusions did not depend on the choice of signature. Results for the large and small signatures are given in the main text and Supplement, respectively. All downstream analyses were conducted with cell types whose average proportions were at least 0.01.

**Figure 1:**
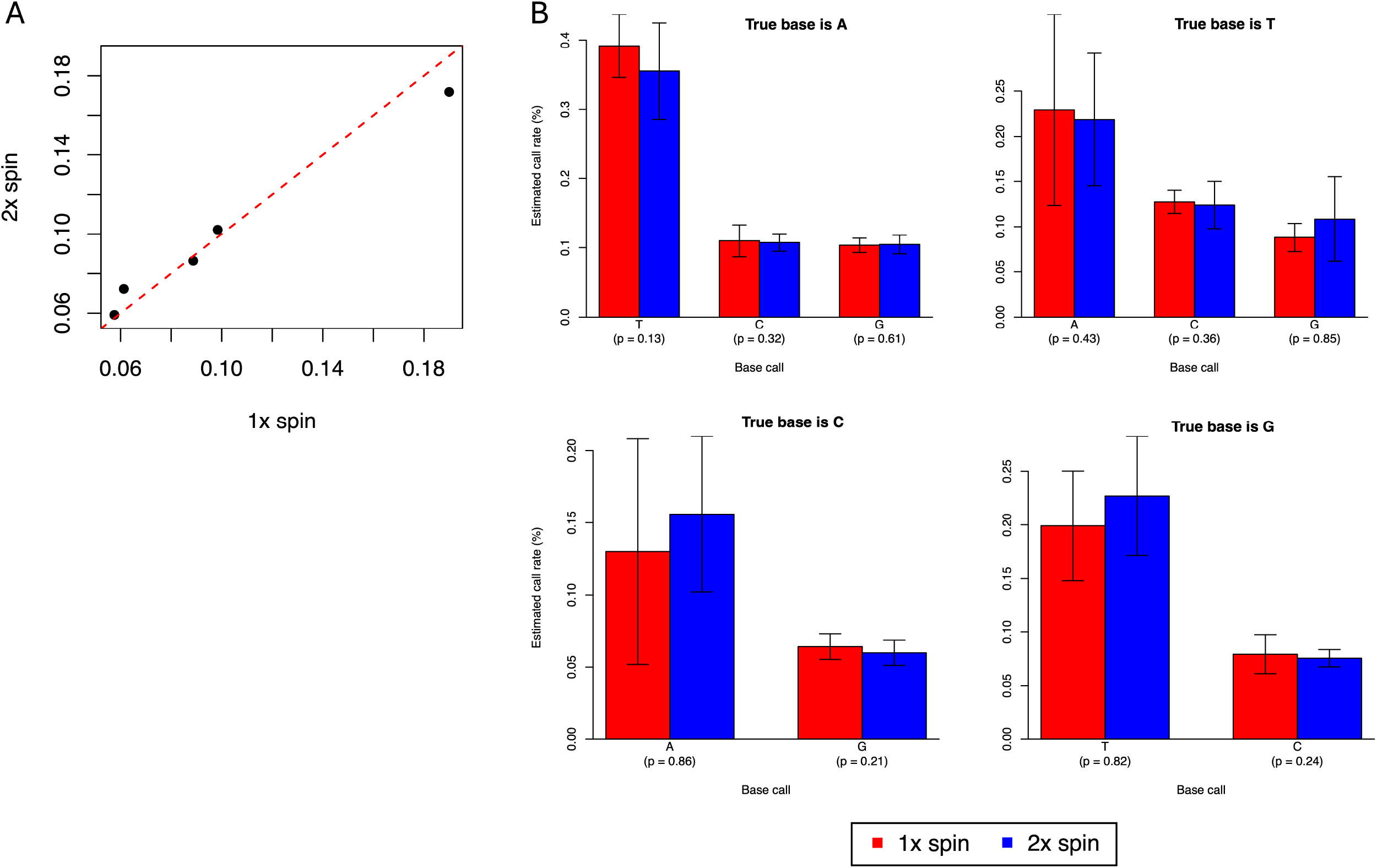
A comparison of cell type proportion estimates in 1x and 2x spin data. Cell types with high, medium, and low proportions are defined based on average proportions. Scatter plots (top) plot estimates for subjects with both 1x and 2x spin measurements. Tables (bottom) report the ICC for each cell type. P-values test the null hypothesis that the ICC is 0. ICCs and p-values were computed using data from all subjects.

We took a model-based approach to determine the impact of the number of spins on proportion estimates. Let *y*_*is*_ be a cell type’s proportion estimate from subject *i* and spin number *s*, where *s* is 1 (single) or 2 (double). Our model was thus defined as *y*_*is*_ =*μ* + *δ*_*i*_ + *∈*_*is*_, where *δ*_*i*_ ∼ *N*(0, *τ*^2^) is subject-specific (biological) variation that is shared across spins, and 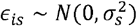 is spin-specific (technical) error. Our hypothesis is that there is no improvement in accuracy for plasma samples that underwent a second spin, i.e., the technical error for one and two spins is the same. To this end, we conducted a likelihood ratio test to evaluate the null hypothesis 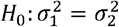 against the alternative 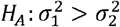 Here, large p-values provide support for *H*_0_, that one and two spins are equally accurate Since we found *τ*^*2*^ was likely non-zero for nearly all cell types, we took the asymptotic distribution for the log-likelihood ratio to be a 50-50 mixture of a point mass at zero and a chi-squared distribution with one degree of freedom. We additionally estimated the intrasubject correlation coefficient (ICC) as the correlation between *y*_*i*1_ and *y*_*i*2_, and tested the null hypothesis that this correlation coefficient was zero against the alternative that it was greater than zero using a likelihood ratio test. Since we had no evidence supporting the hypothesis that 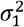 differed from 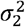 for any cell types, ICC analyses were completed assuming 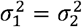.

### Comparing Methylation Rates at Individual CpGs

We performed two parallel analyses to compare the estimates for the methylation rates at individual CpGs derived from single and double-spun data. The first is analogous to our cell type proportion analysis and explicitly examines the hypothesis that methylation rate estimates in single and double-spun data are equally accurate. Let *p*_*is*_ be the methylation rate at a CpG for subject *i* and spin number *s* (1 or 2), where *p*_*is*_ is the probability that the CpG is methylated in that sample. Since *p*_*is*_ is bounded between zero and one, it is difficult to model its variation with Gaussian distributions. Instead, we transformed *p*_*is*_ using the probit function Φ ^− 1^ which is the inverse of the cumulative distribution function for the standard Gaussian distribution and maps the unit interval to the whole real line. We took our model to be *μ*_*is*_= Φ^− 1^ (*p*_*is*_) = *μ* + *δ*_*i*_ + *∈*_*is*_, where, as above, *δ*_*i*_ ∼ *N*(0, *τ*^2^) is biological variance shared by samples and 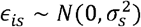 is idiosyncratic and spin number-dependent technical variation. Our goal was to test the null hypothesis 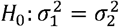 against the alternative 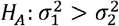 where *H*_0_ implies methylation rates in single and double-spun data are equally accurate and *H*_*A*_ implies single-spun methylation rates are noisier than double-spun rates. Large p-values provide support for *H*_0_. Since CpG read depths are at most finite, *p*_*is*_ is not observed. Instead, we observe 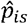, the fraction of times the CpG is observed to be methylated, which is a noisy estimate for *p*_*is*_. Assuming binomial sampling error, an application of the Delta method implies 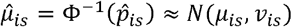, where 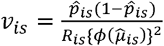. Here *R*_*is*_ CpG’s read depth and *ϕ* is the density function of the standard Gaussian. We used a likelihood ratio test to evaluate *H*_0_ and additionally estimated the ICC as we did above. Since we estimated that the sampling variance *v*_*is*_ significantly exceeded the variance of *μ*_*is*_ when the read depths were small to moderate, we only considered CpGs with median read depths greater than or equal to 40 in this analysis.

In our second analysis, we used all CpGs to compute the correlation between estimated methylation rates for all pairs of methyl-seq libraries. If there is no difference between single and double-spun methylation rates, the correlation between single and double-spun pairs should clearly exceed the correlation between non-pairs. We used a Beta-Binomial-Corrected (BBC) method (see below) to ensure CpGs with low read depths did not obfuscate correlations.

### Beta-Binomial-Corrected Correlation Method

We note that the average methylation rate of a CpG site must be between 0 and 1, matching exactly the support of a Beta distribution. Conditional on the underlying methylation rate of a given CpG site in a sample, the distribution of the observed counts of methylated CpGs from a methyl-seq library follows apparently a Binomial distribution. More importantly, we observed that the histogram of the methylation rates of the CpGs in a methyl-seq library has a distinct U shape, which is very similar to a Beta-Binomial distribution with parameters *α* < 1 and *β*. < 1. Therefore, we used the Beta-Binomial distribution to model the observed methylation rates in methyl-seq libraries.

Consider, for *i*=1,2, two Beta-distributed random variables *Z*_1_ and *Z*_2_ such that *Z*_*i*_ ∼*Beta* (*α*_*i*_, *β*_*i*_) and *Corr*(*Z*_1_,*Z*_2_)=*r*, and two random variables *X*_1_ and *X*_2_ such that *X*_i_ ∼ *Binimial* (*n*_*i*_,*Z*_*i*_),*X*_1_ and *X*_2_ are independent given *Z*_1_,*Z*_2_, where *n*_1_ and *n*_2_ are positive integers. Clearly, *X*_1_ and *X*_2_ both have the marginal Beta-Binomial distribution. Moreover, the correlation between *X*_1_ and *X*_2_ is:

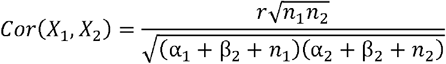

Let *Z*_1_ and *Z*_2_ be the underlying methylation rates of the CpGs in two samples, *X*_1_ and *X*_2_ be the distribution of the observed counts of methylated CpGs for the corresponding two methyl-seq libraries, *n*_1_ and *n*_2_ be the average coverage of the CpGs in the two libraries. It then can be shown that we can estimate *r* = *Cor*(*Z*_1_,*z*_2_) in the following way:

First, estimate the parameters α_*i* +_β_*i*_ and *n*_*i*_ for the two libraries. Specifically, we can estimate α_*i*_ + β_*i*_ from *n*_*i*_ and estimation of 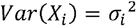 and *E*[*X*_*i*_] = *p*_*i*_ by the method of moments:

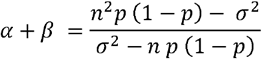

Next, we measure the observed correlation 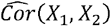 Although the CpG methylation rates are spatially correlated, given the large number of CpGs in each methyl-seq library, this should be a very accurate estimation. Here we ignore the fact that the coverage of the CpGs are not identical within in a library. Finally, the estimated value of *r* is:

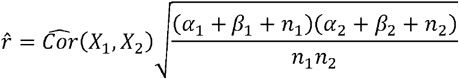

We call this estimated correlation as the Beta-Binomial-Corrected (BBC) correlation.

### cfDNA Fragment Length Comparison

We performed a graphical and statistical comparison of cfDNA fragment lengths between single and double-spun samples as an additional measure of their contrast. Visualizations for read length distributions were produced using the ggplot2 package in R. A Wilcoxon signed-rank test was used to evaluate differences in mean cfDNA fragment lengths between paired single and double-spun samples.

### Genotype-Based Fetal Fraction Estimates

We used base calls at biallelic single nucleotide polymorphisms (SNPs) to estimate the fraction of all fragments originating from the fetus. To ensure estimates were robust to base call errors, we developed a model that accounts for these errors. Briefly, we used 1000 Genomes (Fairley et al., 2020) to identify SNPs with minor allele frequencies ≥0.05 in the CEU superpopulation. We considered CEU because study subjects self-identified as White. We excluded C/T and G/A SNPs because genotypes at these SNPs cannot be inferred due to bisulfite conversion and only used SNPs with at least 5 reads to estimate fetal fraction. We detail our model in the Supplement and use it to estimate both fetal fraction and error rates, i.e., the probability a read is observed to be base *b*_1_ given that the true base is *b*_2_ for all *b*_1_,*b*_2_ ∈{*A,C*,*T*,*G*}. We compared fetal fraction and error rate estimates made in single and double-spun samples.

## RESULTS

### Cellular Proportion Estimates from Single and Double Centrifugation Protocols are Equally Accurate

We sought to determine whether cellular proportion estimates derived from whole genome DNA methylation sequencing data of single-spun maternal plasma cfDNA were as accurate as those from double-spun sample data. Figure 1 gives the concordance between estimates from single and double-spun data, where the scatter plots (top) and corresponding intraclass correlation coefficients (ICCs; bottom) indicate estimates are highly correlated for nearly all cell types.

While these results suggest that the single and double-spun estimates are equally accurate, they are not definitive, as some cell types, such as erythrocyte progenitor and blood macrophage, exhibit modest or weak correlations. One of two hypotheses, *H*_*0*_ and *H*_*A*_, can explain these poor correlations. The null hypothesis, *H*_*0*_, states the proportions of these cell types are intrinsically hard to estimate and will be noisy regardless of the number spins. The alternative hypothesis, *H*_*A*_, states the noise level in single-spun estimates is exceedingly large. Hypothesis *H*_*0*_ occurs when cell type-specific methylation is variable across subjects or if two or more cell types have correlated methylation signature (Huang et al., 2025). If *H*_*0*_ were true, it would imply single and double-spun data are equally accurate. Figure 2A provides a graphical illustration of these two hypotheses, where the first column corresponds *H*_*0*_ and the second to *H*_*A*_. We explicitly tested *H*_*0*_ against *H*_*A*_ (see Methods), where the exceedingly large p-values in Figure 2B indicate there is no evidence to suggest that *H*_*A*_ is true. That is, proportion estimates from single and double-spun data are equally accurate.

**Figure 2:**
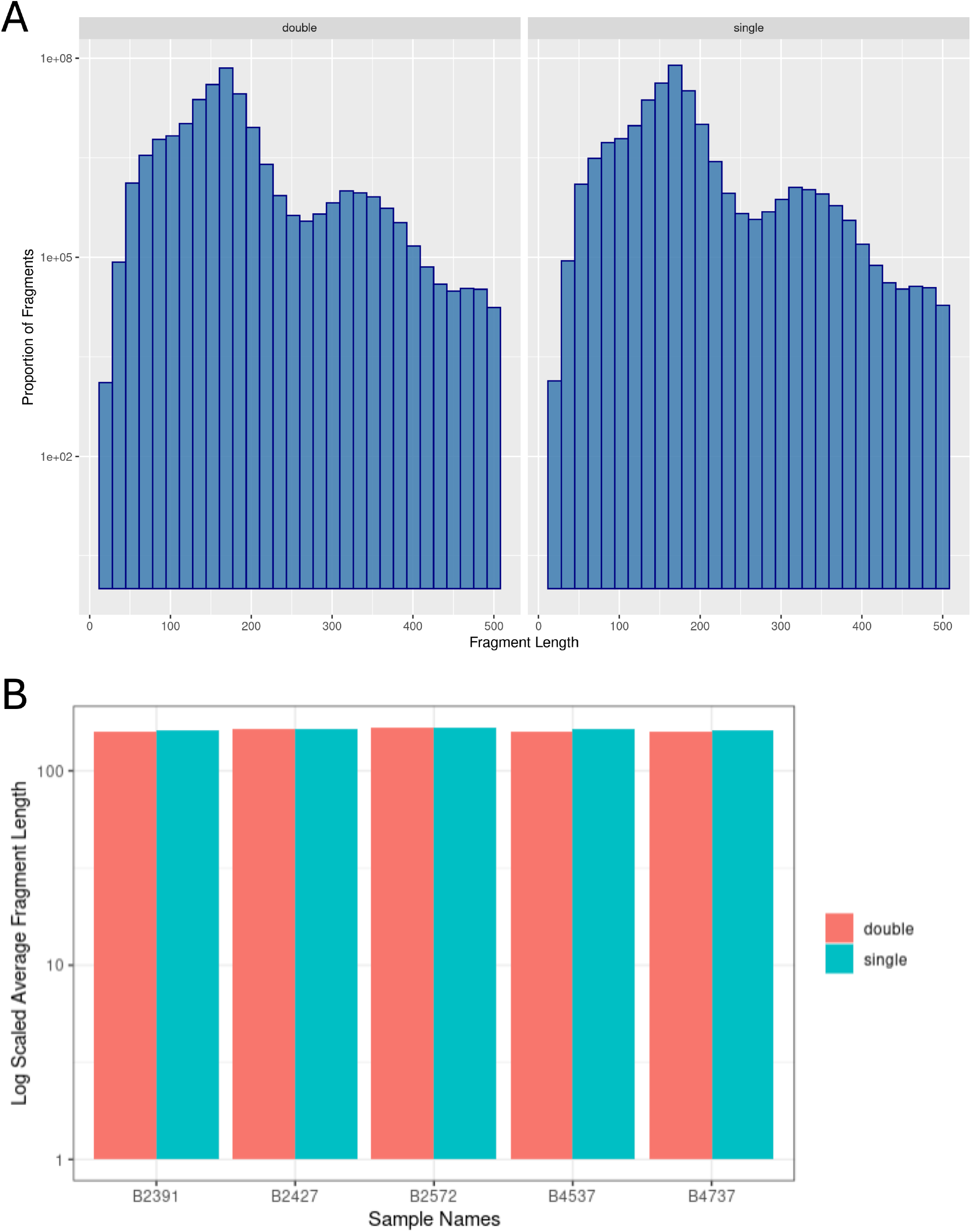
Hypothesis testing to test whether single and double centrifugations give equally accurate cell type proportion estimates. **(A)** Hypothesis table. Columns give the hypotheses and rows the number of spins. Estimates from 1x and 2x spins will be equally accurate if they have the same biological and technical variances (first column); 1x spin estimates will be less accurate if they exhibit greater technical variance than 2x spin estimates (second column). **(B)** P-values for the null hypothesis that 1x and 2x spin data are equally accurate versus the alternative hypothesis that 1x spin estimates are less accurate. Large p-values indicate 1x and 2x spin data are equally accurate.

### DNA Methylation at Single CpG Sites is not Impacted by the Number of Centrifugations

We took a similar approach to the one above to determine whether the fidelity of DNA methylation rates at individual CpGs was not impacted by the number of spins. Due to the relatively small inter-subject variation in rates and our modest sample size, we only considered the 3,253 CpGs with a read depth of at least 40 in this analysis to mitigate the impact of sampling variation (see Methods). Using the same hypotheses considered in Figure 2A, we tested the null hypothesis that single and double-spun data were equally accurate against the alternative that they were not. In line with the cellular proportion results, we found that all 3,253 false discovery rate-adjusted p-values were 1, suggesting that methylation rates from single and double-spun data are equally accurate.

We next investigated the correlation between methylation rates for all pairs of samples (see Methods). Figure 3 gives the results. The dendrogram in the plot shows the hierarchical clustering of the samples based on the correlation matrix. From the heatmap and the clustering, it is clear that the pairs of single-spun and double-spun samples from the same subject have the highest correlations. Figure 4 additionally shows that the density of methylation rates for pairs of single and double-spun samples almost perfectly overlap. These results further suggest that the variation between single vs. double-spun, if any, is far less than the variation between biological replicates.

**Figure 3:**
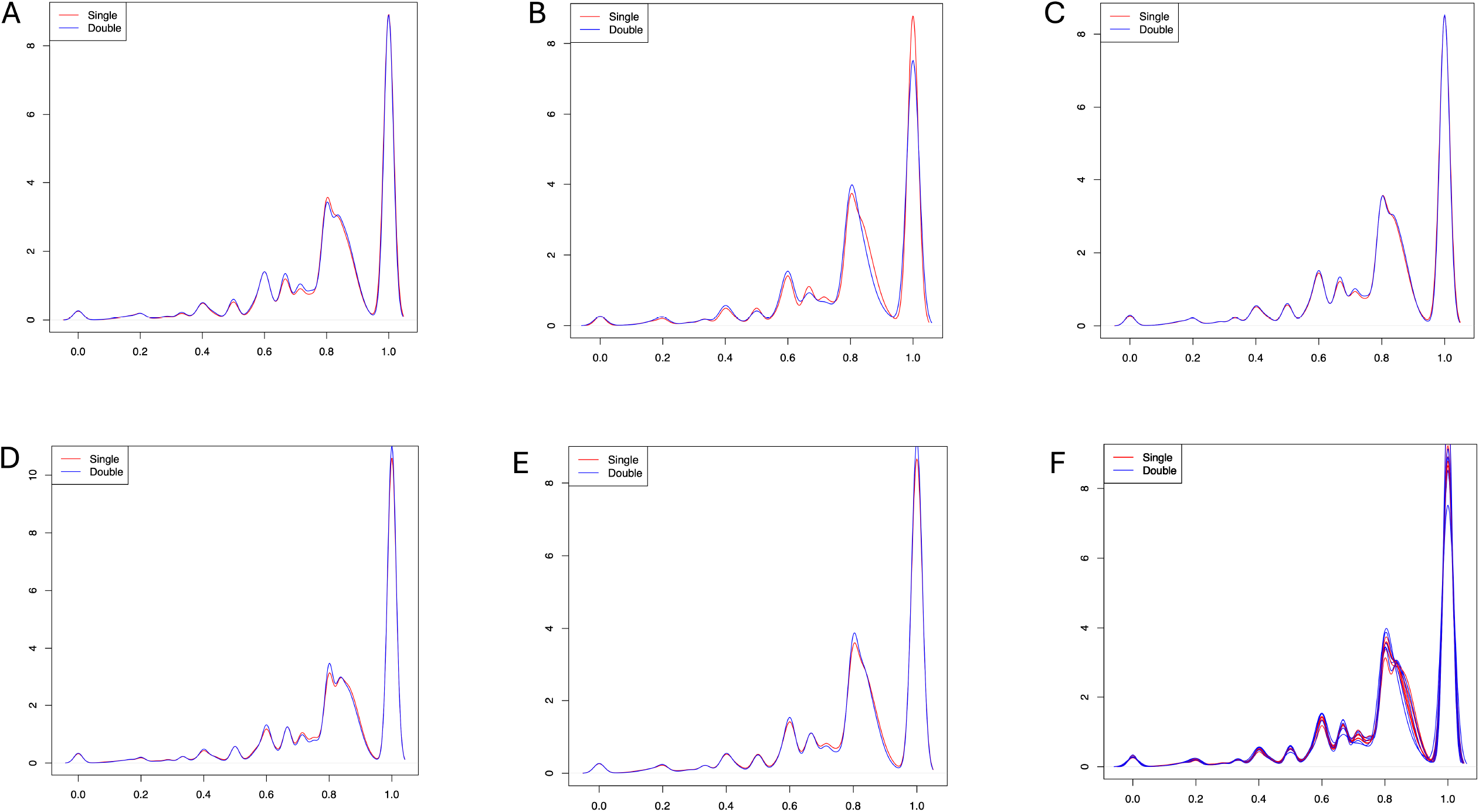
Heatmap for the Beta-Binomial-Corrected correlation matrix for the observed CpG methylation rates of the 11 single and double-spun methyl-seq libraries. Libraries are clustered using the hierarchical clustering algorithm. Libraries with same prefix are from the same patient. Color of the side bars indicates whether a library is single or double-spun.

**Figure 4:**
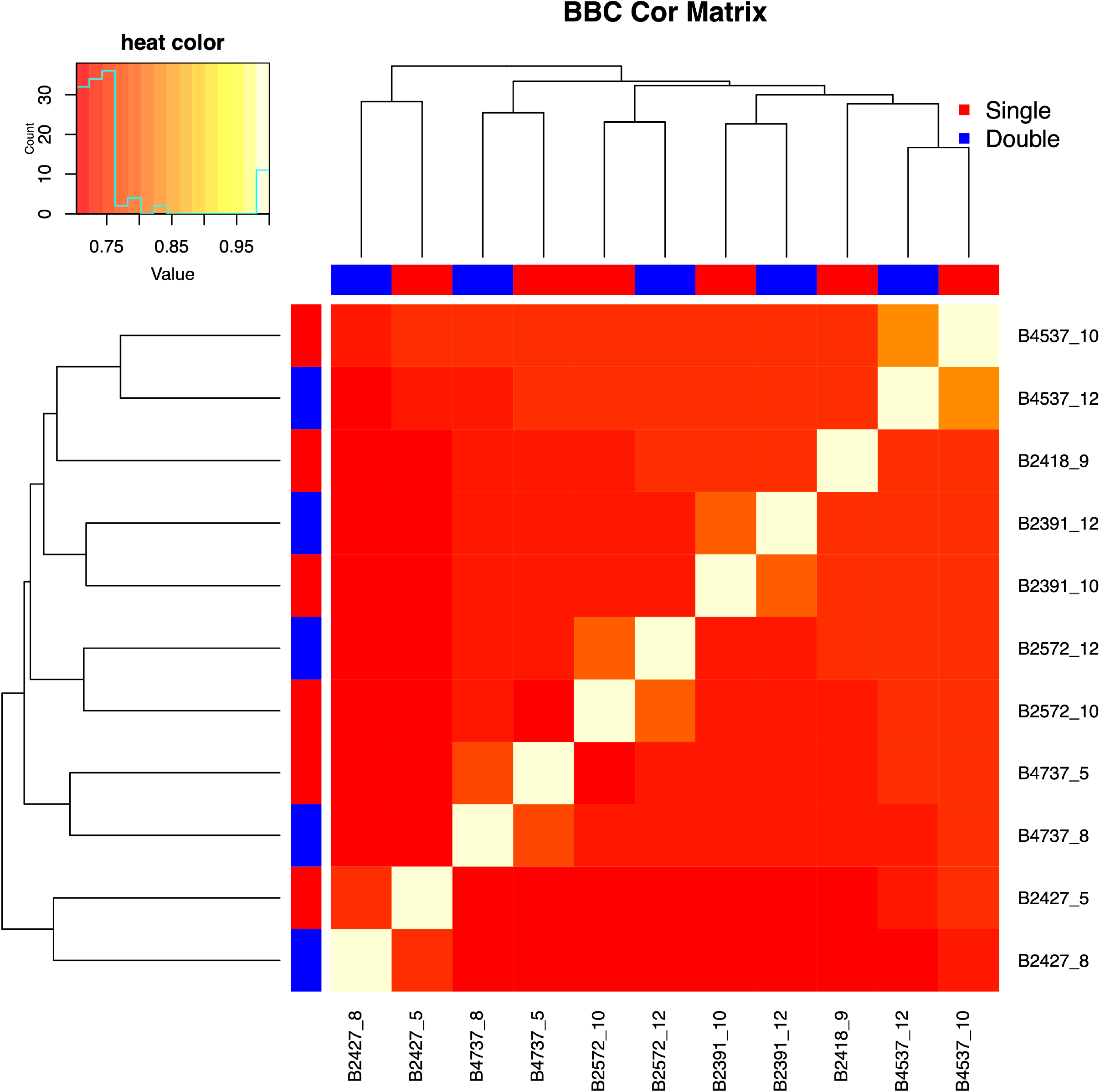
Kernel density estimation (KDE) curves for the distribution of the observed CpG methylation rates. Red curves represent single-spun libraries; blue curves represent double-spun libraries. **A-E:** Paired single/double-spun libraries respectively for patients B2391, B2427, B2572, B4537, B4737. **F:** Combination of A-E.

### Fragment Length Distributions do not Differ Between Single and Double-spun Sample Pairs

cfDNA fragment lengths are an important biomarker. For example, previous investigations have shown that fetal cfDNA fragments are, on average, shorter in length than those of maternal origin (Liang et al., 2018, Qiao et al., 2022). We therefore compared the cfDNA fragment lengths from paired single and double-spun plasma samples. A histogram of fragment lengths reveals that the distribution of fragments for single-spun samples is almost indistinguishable from that for double-spun samples (Figure 5A). Additionally, mean fragment length showed no difference between single and double-spun preparations when comparing across paired samples (Figure 5B; p = 0.4375). We therefore conclude that the proportion of short, fetus derived, fragments is not impacted by a single centrifugation step.

**Figure 5.**
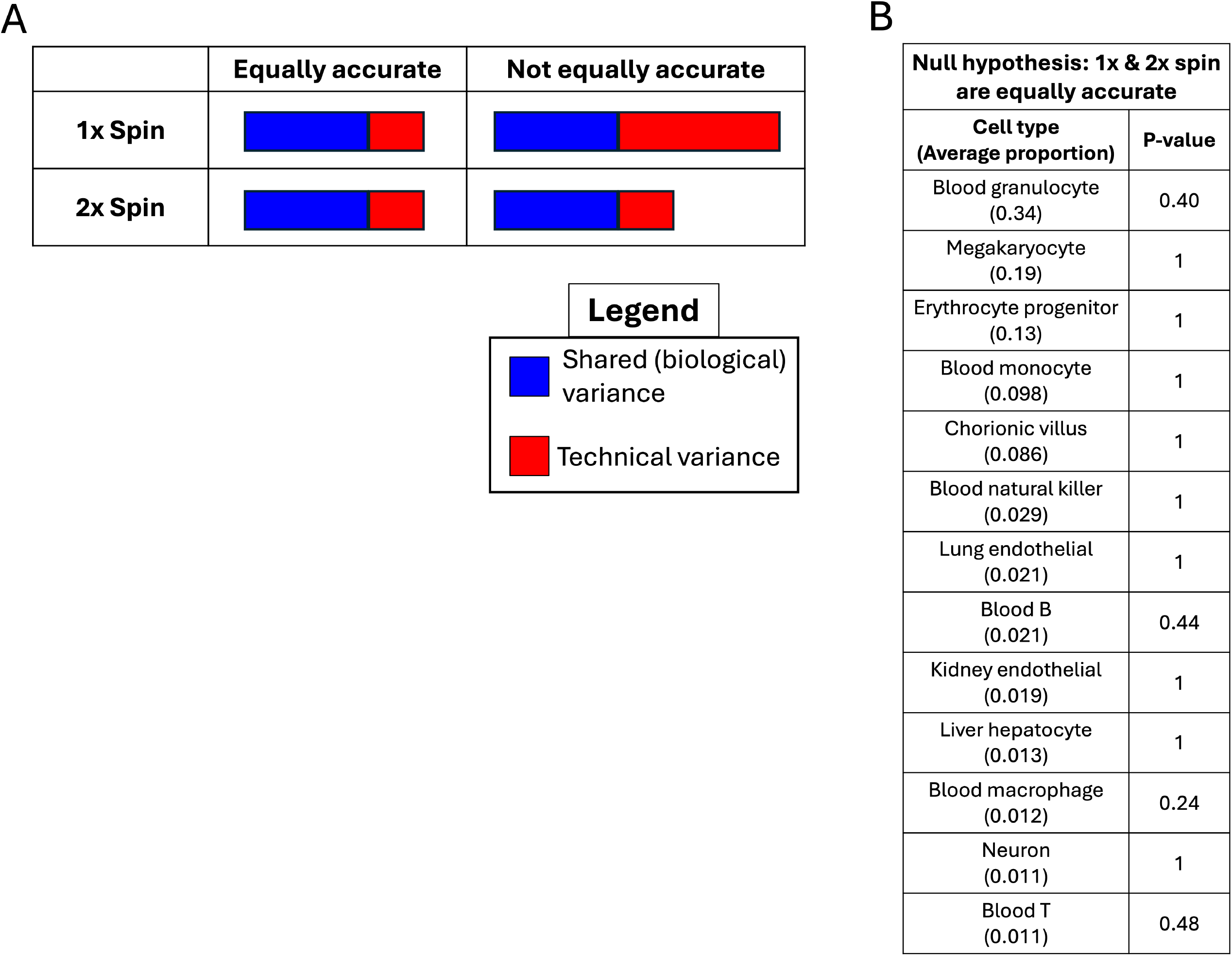
**A:** Histograms for cfDNA fragment lengths across single and double-spun samples. Each distribution is composed of fragments originating from 5 paired plasma samples. Single and double-spun distributions show a high degree of similarity. **Figure 5B:** Mean fragment length across single and double-spun preparation within paired samples. There is no evidence that the median difference in mean fragment length of pairs of single and double-spun samples differs from 0 (p = 0.4375).

### Genotype-Based Fetal Fraction Estimates do not Differ in Single and Double-Spun Samples

Finally, we compared genotype-based estimates for the fraction of all cfDNA fragments originating from the fetus, computed in single and double-spun data. Again, our rationale was that fetal cfDNA fraction would be reduced in single vs. double-spun plasma due to increased presence of high molecular weight DNA from lysed nucleated cells. Figure 6A indicates fetal fraction estimates are virtually identical in single and double-spun samples (ICC=0.99, p-value=1.1×10^−6^). Additionally, base calling error rates at single nucleotide polymorphisms did not depend on the number of spins (Figure 6B), indicating the fidelity of fetal fraction estimates and, more generally, genotype calls were the same in single and double-spun data.

**Figure 6.**
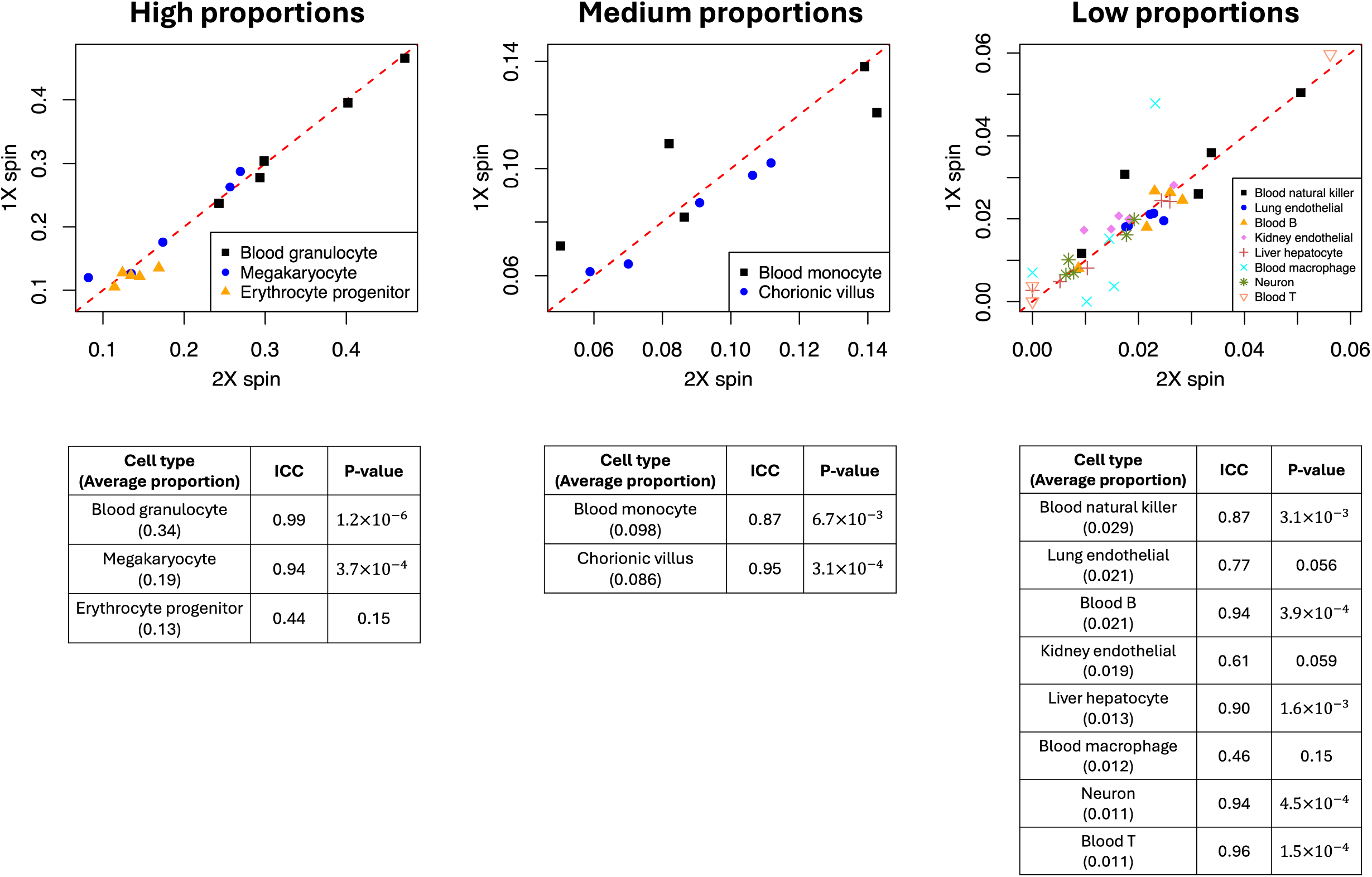
**A:** Genotype-based fetal fraction estimates in single (1x) and double (2x) spun sample pairs. The dashed red line is the line y=x. **Figure 6B:** Estimated base call error rates at single nucleotide polymorphisms used to estimate fetal fraction. P-values (p) test the null hypothesis that the error rate is the same in single and double-spun samples against the alternative that it is greater in single-spun samples. Large p-values suggest no difference between the number of spins. Error bars give 95% confidence intervals. Errors due to deamination (C -> T and G -> A) are not plotted.

## DISCUSSION

Standardized preanalytical processing of peripheral blood samples is widely regarded as a critical element of accuracy and reproducibility for cfDNA-based liquid biopsy assays. It is broadly accepted that the removal of high-molecular-weight genomic DNA from lysed nuclei is critical to maximize cfDNA fraction in DNA extracted from plasma (Jahr et al., 2001, Lo et al., 1997). Standard practice consists of an initial low-speed centrifugation step to separate plasma from other components, followed by a second centrifugation step at much higher speed to ensure maximal removal of nucleated cells. This approach has been widely adopted across oncology, prenatal testing, and transplantation (Chiu et al., 2001, El Messaoudi et al., 2013).

In this study we performed a systematic and quantitative evaluation of whether a second high-speed centrifugation step confers measurable analytical benefits when plasma is initially separated using a single low-speed spin under controlled conditions. By analyzing paired single- and double-spun plasma aliquots derived from the same blood draws, we were able to investigate the effect of the number of centrifugation steps without the confounder of biological variability. In all contexts, including the DNA methylation-based evaluation of cell-type proportion via deconvolution, single CpG-level methylation rate estimation, fragment length analysis, and genotype-based fetal fraction inference, we found no evidence that double centrifugation improved accuracy or reduced technical noise relative to single centrifugation.

Evidence that a second centrifugation step is required to minimize nuclear DNA contamination (and effective dilution) of cfDNA was originally obtained via the use of polymerase chain reaction (PCR)-based methods. This is important because PCR methods are unable to directly distinguish differences in the fragment lengths of template DNA from which amplification products are derived. Thus, cfDNA and nuclear DNA fragments from the same genomic loci are indistinguishable and cfDNA fraction estimations must be made by targeting sequences present only in the cfDNA, such as the Y chromosome of a male fetus in the context of pregnancy (Chiu et al., 2001).

Molecular methods have evolved considerably since preanalytical protocols for cfDNA preparation were established. This is significant because cfDNA-compatible sequencing library preparation workflows across platforms are inherently permissive of short fragments and operationally inefficient with longer molecules. While modern genomic library preparation methods don’t explicitly remove long genomic DNA, their chemistry and workflow design are optimized to efficiently process DNA fragments in the cfDNA optimal size range of ∼150-300bp. This selection is driven at multiple points in the protocol, beginning with end-repair and adapter ligation steps, where shorter fragments are favorably selected over long genomic DNA due to their low steric hindrance and improved end accessibility. Favorable selection of shorter fragments proceeds further during wash/purification steps during which SPRI bead ratios favor shorter fragments due to their increased binding and elution efficiencies. Subsequent PCR amplification of the libraries further reinforces this bias as short fragments preferentially amplify more efficiently and reproducibly.

Cell-type proportion estimation using cfDNA methylation signatures provides a particularly sensitive test of genomic DNA contamination, as leukocyte-derived nuclear DNA would be expected to inflate contributions from nucleated hematopoietic populations and distort inferred tissue mixtures. Using both large and small reference signatures derived from genome-wide methylation atlases (Loyfer et al., 2023), we observed strong concordance between single- and double-spun estimates across nearly all cell types (Figure 1). Scatter plots and corresponding intraclass correlation coefficients demonstrate that paired estimates cluster tightly along the identity line, indicating high agreement between centrifugation protocols. For the few cell populations exhibiting weaker correlations, we explicitly tested whether this variability reflected excess technical noise in single-spun plasma. The results strongly supported the null hypothesis that technical variance did not differ between single- and double-spun samples (Figure 2), indicating that these weaker correlations arise from intrinsic biological heterogeneity or overlapping methylation signatures rather than from inadequate plasma clarification. These findings are consistent with prior work showing that deconvolution accuracy is often limited by reference collinearity and inter-individual epigenetic variability rather than sequencing depth or library preparation artifacts (Caggiano et al., 2021, Huang et al., 2025).

At the level of individual CpG sites, we again found no evidence that methylation rate estimates were degraded in single-spun plasma. By explicitly modeling biological variance, technical variance, and binomial sampling noise, we tested whether single-spin processing introduced additional error. Across all CpGs meeting conservative read-depth thresholds, false discovery rate-adjusted likelihood ratio tests uniformly supported the null hypothesis of equal accuracy between data from single- and double-spun plasma.

Correlation-based analyses using a beta-binomial framework further demonstrated that paired single- and double-spun libraries from the same individual were consistently more similar to each other than to any other sample. Hierarchical clustering based on corrected correlations grouped paired samples together with high confidence (Figure 3), and the genome-wide distribution of CpG methylation rates showed near-complete overlap between single- and double-spun libraries (Figure 4). Together, these results indicate that centrifugation number contributes negligibly to CpG-level methylation variability relative to biological differences between individuals.

cfDNA in maternal plasma originating from the fetus/placenta is known to be of overall shorter length than fragments of maternal origin (Qiao et al., 2019). Therefore, the increased presence of maternal genomic DNA caused by inadequate removal of nucleated blood cells would be expected to result in a shift towards longer cfDNA fragment distributions. However, we found that fragment length distributions from paired single- and double-spun plasma samples were nearly indistinguishable (Figure 5A), with no systematic differences in the relative abundance of long fragments or short-fragments. Mean fragment lengths did not differ between protocols when compared within subjects (Figure 5B), further indicating that residual cellular material after a single low-speed spin does not measurably contribute to contaminating DNA under the conditions tested. Similarly, genotype-based fetal fraction estimates were virtually identical between single- and double-spun samples. Because fetal fraction is particularly sensitive to background maternal genomic DNA, even modest leukocyte lysis would be expected to reduce inferred fetal contribution and increase genotype calling noise (Chan et al., 2005). Instead, fetal fraction estimates showed near-perfect concordance across centrifugation protocols (Figure 6A), and base-calling accuracies at informative SNPs were similarly indistinguishable (Figure 6B). These results provide strong evidence that single-spun plasma is suitable for allele-specific cfDNA analyses, including non-invasive prenatal testing workflows.

Our findings indicate that, when blood is collected in EDTA tubes and plasma is separated promptly using careful aspiration that avoids disturbance of the buffy coat, a single low-speed centrifugation step is sufficient to support high-fidelity cfDNA methylation, fragmentomic, and genotyping analyses.

Importantly, these results do not suggest that double centrifugation is unnecessary in all circumstances. Rather, they indicate that the analytical benefits of an additional high-speed centrifugation step may be negligible under controlled processing conditions in which leukocyte lysis is minimal. In routine clinical practice, double centrifugation may remain advisable when sample handling is delayed, when plasma aspiration is performed under suboptimal conditions, or when the integrity of the cellular fraction is uncertain. From a practical perspective, these findings suggest that plasma collections generated using single-spin protocols may be suitable for genome-wide cfDNA analyses. This greatly increases the potential for retrospective analysis of stored plasma samples that have not been collected and processed specifically for cfDNA studies and suggests that clinical laboratory workflows could be shortened by removing the second spin.

This study has several limitations. First, the number of subjects analyzed was modest, and all samples were derived from early-pregnancy maternal plasma; therefore, our findings may not generalize to other clinical contexts such as oncology or transplant monitoring, where cfDNA abundance, composition, and background leukocyte burden can differ substantially. Second, all samples were collected in EDTA tubes and processed promptly under controlled conditions with careful plasma aspiration, and thus these conclusions may not apply to specimens collected in alternative tube types, subjected to delayed processing, or handled under less controlled pre-analytical conditions. In addition, while the paired design provides strong sensitivity to detect large technical differences between single- and double-centrifugation protocols, the modest sample size limits power to detect more subtle effects. Accordingly, the absence of statistically significant differences should not be interpreted as evidence that such differences cannot occur. Rare leukocyte lysis events, ultra-low allele fraction signals, or edge-case pre-analytical failures may introduce effects that are not captured in a small, well controlled cohort. Finally, we did not directly assess downstream clinical assay performance, such as aneuploidy detection or somatic mutation calling at very low variant allele fractions, nor did we examine the impact of additional variables such as repeated freeze-thaw cycling or long-term storage. Despite these limitations, the paired experimental design and concordant results across multiple independent cfDNA metrics provide strong evidence that, under the conditions tested, a second centrifugation step does not measurably improve analytical fidelity.

Collectively, these findings challenge the prevailing assumption that double centrifugation is absolutely required for reliable cfDNA analysis in peripheral blood plasma. While additional centrifugation may provide a safeguard against suboptimal handling or delayed processing, our results demonstrate that a single low-speed centrifugation step is sufficient for high-resolution cfDNA methylation, fragment-based, and genotyping assays. This suggests that existing biorepositories containing single-spun plasma samples can be used for liquid biopsy research and assay development, which will accelerate discovery and clinical translation across multiple liquid biopsy applications.

