## Supplemental Information for "Re-evaluating the Need for Double Centrifugation in Plasma Cell-Free DNA Analysis"

**SUPPLEMENTAL MATERIAL**

Our goal is to estimate $\pi$, the fraction of paternally derived fragments, where fetal fraction is $2\pi$. To describe our model, let $r_{g}$ be the read depth at SNP $g$, $y_{gb}$ be the number of reads that were called as base $b=A,C,T,\mathrm{or} G$, and $\boldsymbol{y}_{g}$ to be the corresponding length-4 vector of base calls. We condition on $r_{g}$ in all below probability models, though we exclude it from conditioning statements to simplify notation. To account for base-calling errors, define $\epsilon_{b_{1}b_{2}}$ to be the probability a read is observed to be base $b_{1}$ given that the true base is $b_{2}$. Let $\boldsymbol{\epsilon}$ be the corresponding 4x4 error matrix. Then conditional on maternal and paternal genotypes $M_{g}$ and $P_{g}$ at SNP $g$, we assumed $\boldsymbol{y}_{g}$ was multinomial with probability vector $\boldsymbol{p}_{g}$ determined by $\pi$, $\boldsymbol{\epsilon}$, $M_{g}$, and $P_{g}$:

$$\boldsymbol{y}_{g}\mid\left( M_{g},P_{g}, \pi,\boldsymbol{\epsilon} \right)\sim\mathrm{Mult}\left( r_{g},\boldsymbol{p}_{g}\left( \pi,\epsilon,M_{g},P_{g} \right) \right).$$

As there are six possible genotypes (three for the mother times two for the father), $\boldsymbol{p}_{g}\left( \pi,\epsilon,M_{g},P_{g} \right)$ takes one of six values, which are given below assuming the SNP’s alleles are $A$ and $T$:

1. $M_{g}=AA, P_{g}=A$. $p_{gA}=\epsilon_{AA}$; $p_{gC}=\epsilon_{CA}$; $p_{gT}=\epsilon_{TA}$; $p_{gG}=\epsilon_{GA}$
2. $M_{g}=AT, P_{g}=A$. $p_{gA}=\frac{\left( 1+\pi\right)}{2}\epsilon_{AA}+\frac{\left( 1-\pi\right)}{2}\epsilon_{AT}$; $p_{gC}=\frac{\left( 1+\pi\right)}{2}\epsilon_{CA}+\frac{\left( 1-\pi\right)}{2}\epsilon_{CT}$; $p_{gT}=\frac{\left( 1+\pi\right)}{2}\epsilon_{TA}+\frac{\left( 1-\pi\right)}{2}\epsilon_{TT}$; $p_{gG}=\frac{\left( 1+\pi\right)}{2}\epsilon_{GA}+\frac{\left( 1-\pi\right)}{2}\epsilon_{GT}$
3. $M_{g}=TT, P_{g}=A$. $p_{gA}=\pi\epsilon_{AA}+(1-\pi)\epsilon_{AT}$; $p_{gC}=\pi\epsilon_{CA}+(1-\pi)\epsilon_{CT}$; $p_{gT}=\pi\epsilon_{TA}+(1-\pi)\epsilon_{TT}$; $p_{gG}=\pi\epsilon_{GA}+(1-\pi)\epsilon_{GT}$
4. $M_{g}=AA, P_{g}=T$. $p_{gA}=(1-\pi)\epsilon_{AA}+\pi\epsilon_{AT}$; $p_{gC}=(1-\pi)\epsilon_{CA}+\pi\epsilon_{CT}$; $p_{gT}=(1-\pi)\epsilon_{TA}+\pi\epsilon_{TT}$; $p_{gG}=(1-\pi)\epsilon_{GA}+\pi\epsilon_{GT}$
5. $M_{g}=AT, P_{g}=T$. $p_{gA}=\frac{\left( 1-\pi\right)}{2}\epsilon_{AA}+\frac{\left( 1+\pi\right)}{2}\epsilon_{AT}$; $p_{gC}=\frac{\left( 1-\pi\right)}{2}\epsilon_{CA}+\frac{\left( 1+\pi\right)}{2}\epsilon_{CT}$; $p_{gT}=\frac{\left( 1-\pi\right)}{2}\epsilon_{TA}+\frac{\left( 1+\pi\right)}{2}\epsilon_{TT}$; $p_{gG}=\frac{\left( 1-\pi\right)}{2}\epsilon_{GA}+\frac{\left( 1+\pi\right)}{2}\epsilon_{GT}$
6. $M_{g}=TT, P_{g}=T$. $p_{gA}=\epsilon_{AT}$; $p_{gC}=\epsilon_{CT}$; $p_{gT}=\epsilon_{TT}$; $p_{gG}=\epsilon_{GT}$

Values for other SNPs are analogous. The full data log-likelihood can then be expressed as

$$\mathcal{l}\left( \pi,\boldsymbol{\epsilon} \right)=\sum_{g} \log\left\{ \sum_{M_{g},P_{g}} \Pr\left( \boldsymbol{y}_{g} \mid M_{g},P_{g},\pi,\boldsymbol{\epsilon} \right)\Pr\left( M_{g} \right)\Pr(P_{g}) \right\},$$

where the first sum is taken over all SNPs $g$ and the second sum is over all six genotype combinations. The priors on genotypes $\Pr\left( M_{g} \right)$ and $\Pr(P_{g})$ are computed assuming Hardy-Weinberg equilibrium using minor allele frequencies derived from the 1000 Genomes CEU population. Estimates for $\pi$ and $\boldsymbol{\epsilon}$ were derived by maximizing $\mathcal{l}\left( \pi,\boldsymbol{\epsilon} \right)$ under the constraint that $\pi\leq0.5$ (i.e., fetal fraction is $\leq1$) and the entries of $\boldsymbol{\epsilon}$ are $\geq0$ and its columns add up to 1.
